# Marker-based CRISPR screening reveals a MED12-p63 interaction that activates basal identity in pancreatic ductal adenocarcinoma

**DOI:** 10.1101/2023.10.24.563848

**Authors:** Diogo Maia-Silva, Allison C. Schier, Damianos Skopelitis, Vahag Kechejian, Aktan Alpsoy, Jynelle Liverpool, Dylan J. Taatjes, Christopher R. Vakoc

## Abstract

The presence of basal lineage characteristics signifies hyper-aggressive human adenocarcinomas of the breast, bladder, and pancreas. However, the biochemical mechanisms that maintain this aberrant cell state are poorly understood. Here we performed marker-based genetic screens in search of factors needed to maintain basal identity in pancreatic ductal adenocarcinoma (PDAC). This approach revealed MED12 as a powerful regulator of the basal cell state in this disease. Using biochemical reconstitution and epigenomics, we show that MED12 carries out this function by bridging the transcription factor p63, a known master regulator of the basal lineage, with the Mediator complex to activate lineage-specific enhancer elements. Consistent with this finding, the growth of basal-like PDAC is hypersensitive to MED12 loss when compared to classical PDAC. Taken together, our comprehensive genetic screens have revealed a biochemical interaction that sustains basal identity in human cancer, which could serve as a target for tumor lineage-directed therapeutics.

---

Cellular identity is commonly dysregulated in human cancer^1^. As a prominent example, human adenocarcinomas, which are tumors of the glandular epithelial lineage, can acquire characteristics of basal (also known as squamous) epithelial cells during disease progression^2^. This process is most evident in adenocarcinomas of the breast^3,4^, bladder^5,6^, and pancreas^7–10;^ tumors in which expression of basal lineage markers (*e.g. TP63* and *KRT5*) portends an inferior clinical outcome. Emerging evidence in human lung adenocarcinoma also highlights the acquisition of basal characteristics as a means of bypassing EGFR^11^ or KRAS^12^ -targeting therapeutics. While the clinical significance of basal-like adenocarcinomas has become clear in recent years, the biochemical mechanisms that drive this aberrant cell fate remain largely unknown.

One critical master regulator of the basal lineage in normal^13^ and neoplastic contexts^14^ is the transcription factor (TF) p63, the protein product of the *TP63* gene. While largely undetectable in the normal human and mouse pancreas^15,16^, a subset of pancreatic ductal adenocarcinomas (PDAC) acquires p63 expression in close association with a basal-like transcriptome and inferior patient outcomes^9,10,17–19^. We and others previously demonstrated the necessity and sufficiency of p63 to endow PDAC cells with basal lineage characteristics^18,20^, which in turn leads to enhanced cell motility and invasion^18^, stromal inflammation^21^, chemotherapy resistance^22^, and a powerful p63 dependency for cell viability and proliferation^18,21,23^. Despite this potent transcriptional activation function seen *in vivo*, the variant of p63 expressed in basal-like PDAC is the ΔN isoform that lacks its critical N-terminal activation domain^9,18^. The biochemical mechanism by which the ΔN isoform of p63 (hereafter referred to as p63 for simplicity) activates basal identity in PDAC has yet to be defined.

The Mediator complex is a multi-subunit transcriptional coactivator required for most RNA Polymerase II-dependent transcription in eukaryotes^29^. This general transcriptional role depends on the core of Mediator, which is composed of twenty-six subunits that lack any known enzymatic activity. However, a reversibly attached four subunit Mediator kinase module (MKM), comprised of MED12 or MED12L, MED13 or MED13L, Cyclin C, and CDK8 or CDK19, contributes to transcription through phosphorylation of protein substrates^24^. Unlike the broad requirements for core Mediator in transcription, the MKM performs specialized transcriptional functions in a context-specific manner^24–26^.

To complement prior hypothesis-oriented studies of basal-like PDAC^18,20^, we set out to develop a marker-based CRISPR screening method capable of revealing all genes needed to maintain basal identity in this disease (**Fig. 1a**). Our approach relies on intracellular fluorescent activated cell sorting (FACS) staining of p63 and KRT5, which are validated diagnostic markers that distinguish basal-like tumors from classical adenocarcinomas^9,17,27^. After optimizing the staining conditions and the quantitative resolution of the assay (**Fig. 1b, Extended Fig. 1a**), we performed genome-wide CRISPR interference (CRISPRi) screens^28^ in three basal-like PDAC cell line models (T3M4, KLM1, and BxPC3), measuring effects of >20,000 genetic perturbations on KRT5 and p63 expression (**Fig. 1c, Supplementary Table 1**). In all six screens, the outlier performance of *KRT5* and *TP63* sgRNAs supported the overall quality of these datasets (**Fig. 1c**). Since p63 directly activates *KRT5* (**Extended Fig. 1b**) and its own expression at the *TP63* locus^29,30^, we reasoned that any additional outlier hits identified across these screens might represent factors that cooperate biochemically with p63 to activate basal lineage features.

**Fig. 1.**
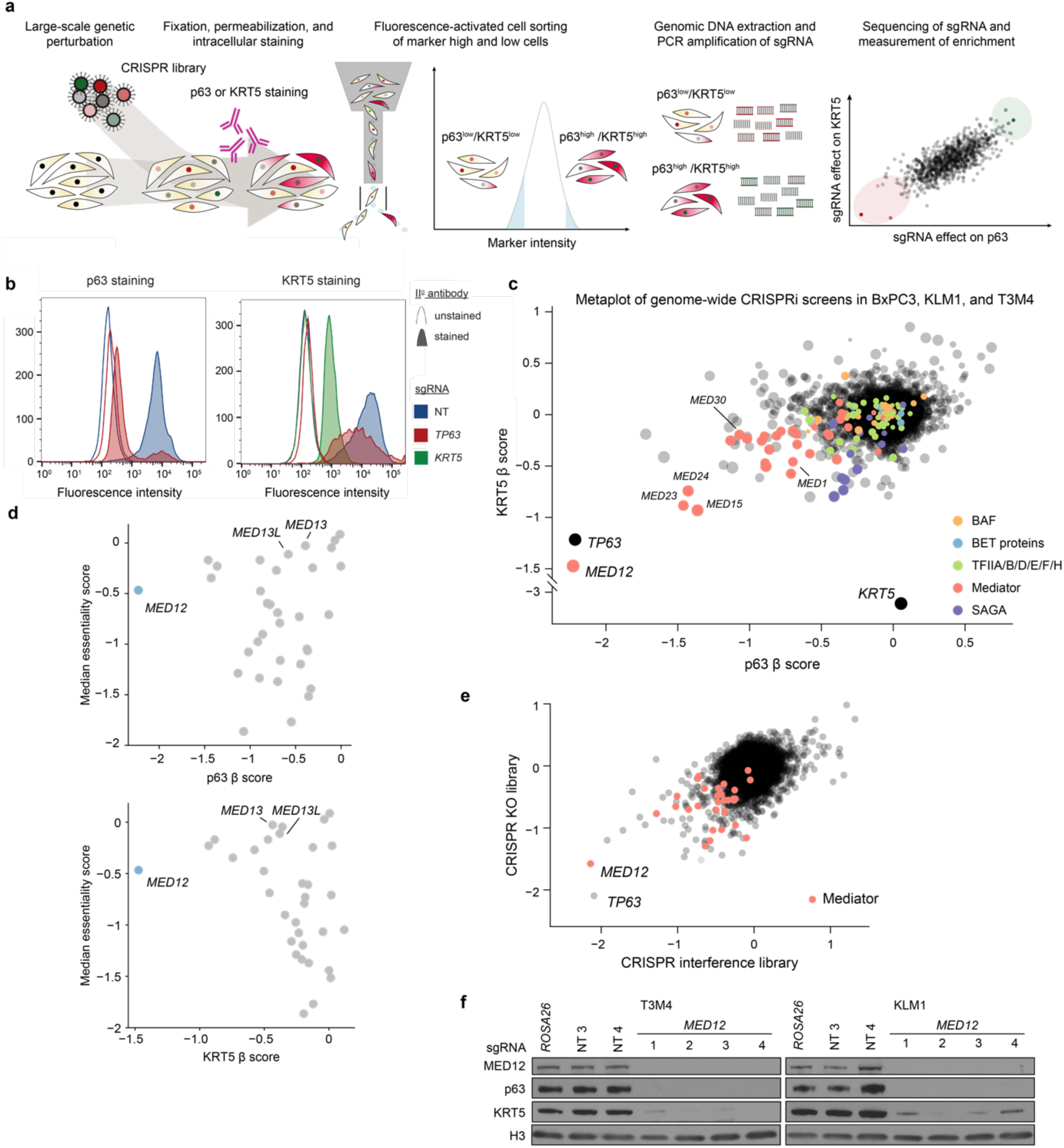
Intracellular FACS-based genome-wide CRISPR screens uncover *MED12* as a critical regulator of basal lineage identity in PDAC. **a,** Diagram illustrating the workflow of KRT5 or p63 genome-wide reporter screens. **b,** Flow cytometry staining profiles for CRISPRi-mediated *TP63* or *KRT5* knockdown in KLM1 cells. Secondary staining with AF647-conjugated anti-rabbit antibody (area-filled curves) or unstained controls (outline-only curves) show the signal distribution of p63-(left) or KRT5-(right) stained cells upon gene knockdown. A minimum of 10,000 events were collected and plotted for each sample. **c,** Metaplot of p63- and KRT5-based reporter genome-wide CRISPRi screen results in 3 independent cell lines (KLM1, T3M4 and BxPC3). Average β scores of p63 and KRT5 screens are plotted such that each dot represents one promoter-defined gene. The size of each dot is proportional to the inverse of the standard deviation of the β values across cell lines. Important mammalian general transcriptional regulators are highlighted according to the legend. β scores were calculated using MAGeCK with the MLE option, with negative β scores denoting enrichment in the marker^low^ population. **d,** Scatterplot of the average β scores in the p63 (top) or KRT5 (bottom) reporter screens across KLM1, T3M4, and BxPC3, and the median CERES cancer cell line essentiality score from the Cancer Dependency Map. **e,** Scatterplot of p63-based reporter screens β scores using genome-wide CRISPRi or CRISPR knockout libraries in KLM1 cells. Genes belonging to the Mediator complex are highlighted in red. **f,** Western blot of whole cell lysates at day 6 post-infection with lentiviral CRISPR knockout sgRNA targeting *MED12* or negative control sgRNAs in T3M4 and KLM1 cells.

All six of our genetic screens independently nominated *MED12* as encoding a top genetic requirement for p63 and KRT5 expression in basal-like PDAC (**Supplementary Table 1**). Notably, the *MED12* requirement for p63 and KRT5 expression exceeded that of all other Mediator subunits and also exceeded the requirement for other general transcriptional machineries (*e.g.* TFIID, p300, BAF complex, BET proteins) (**Fig. 1c**). While most of the core subunits of Mediator are pan-essential dependencies across all cancer cell lines, *MED12* exhibits a high degree of cell line selectivity in its essentiality requirement (**Fig. 1d**). To validate our results, we repeated our marker-based screen using a genome-wide CRISPR knockout sgRNA library^31^, which also recovered *TP63* and *MED12* as outlier requirements for basal lineage identity (**Fig. 1e**). We cloned and tested individual sgRNAs targeting *MED12*, and validated potent downregulation of MED12, p63, and KRT5 at the protein level via western blotting analysis in two different basal-like PDAC models (**Fig. 1f**). Collectively, our screening results provided a strong rationale to investigate MED12 as a regulator of basal identity in PDAC.

We next performed RNA-seq analysis following CRISPR-based targeting of *MED12* and *TP63* in T3M4 and KLM1 cells, which led to a broad suppression of clinically-defined basal lineage signatures^10^ (**Fig. 2a,b, Extended Fig. 2a, Supplementary Table 2**). *MED12* knockout also significantly suppressed a core set of direct p63 target genes in PDAC (**Fig. 2b, Extended Fig. 2a**), which we defined previously^18^. As a control for specificity, knockout of other essential genes (e.g. *CDK1*, *SUPT20H*, and *PCNA*) failed to suppress basal and p63 target gene signatures (**Fig 2c, Supplementary Table 2**). We next sought to distinguish whether MED12 directly activates the basal lineage transcriptome or indirectly supports basal identity by activating *TP63* transcription. Results from RT-qPCR timecourse measurements and from p63 cDNA rescue assays indicated that MED12 functions to directly activate a large program of basal lineage genes, which includes *TP63* (**Fig. 2c,d, Extended Fig. 2b**). These findings indicate that p63 and MED12 are each required to activate the basal lineage signature in PDAC.

**Fig. 2.**
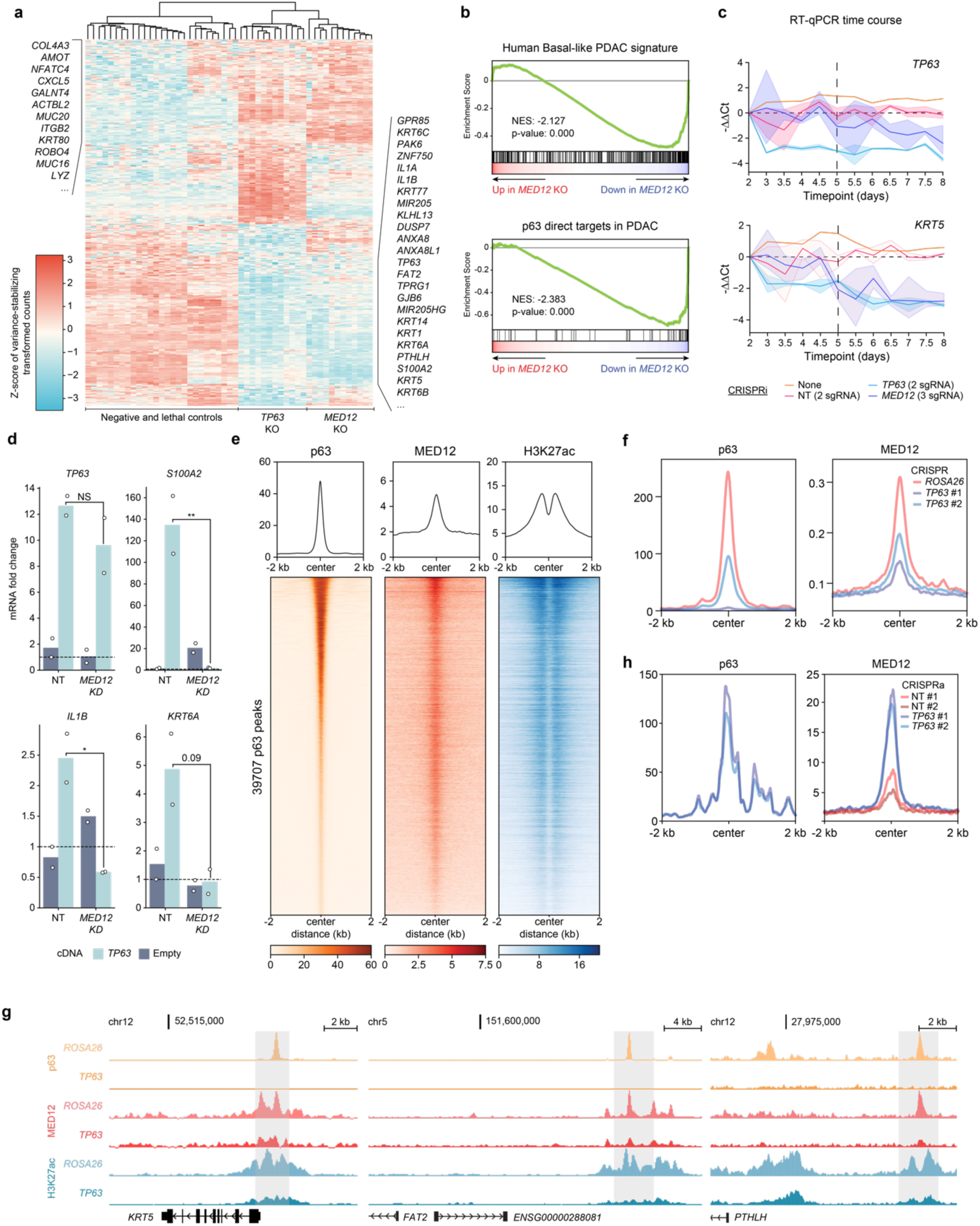
p63 recruits MED12 to chromatin to coactivate the basal transcriptional signature. **a,** Heatmap of z-scored variance-stabilizing transformed gene counts of top 250 overexpressed and downregulated genes upon *TP63* knockout in T3M4 cells. Non-targeting sgRNAs or knockout of *ROSA26* (negative controls); *CDK1*, *PCNA*, or *RPA3* (lethal controls); *TP63*; or *MED12* were performed with two to four different sgRNAs and three biological replicates each. Each column corresponds to an individual RNA-seq sample, and each row displays the z-score of variance-stabilized transformed counts (calculated with DESeq2) across samples. Samples are clustered by Euclidean distance according to the dendrogram on top of the figure. Genes associated with basal (right) and classical (left) PDAC are labeled on the side. **b,** Representative GSEA plots of T3M4 *MED12* knockout using gene signatures derived from human basal PDAC tumors (Chan-Seng-Yue et al. 2020) and direct p63 gene targets in PDAC (Somerville et al. 2018). The plots were generated and the normalized enrichment scores (NES) and statistics were calculated using DESeq2-derived log2FC values of *MED12* vs *ROSA26* KO in GSEApy. Three biological replicates were used for each sample. Complete GSEA analysis for all the sgRNA tested can be found in Supplementary Table 2. **c,** Time-course RT-qPCR of *TP63* and its target gene *KRT5* after lentiviral infection with CRISPRi sgRNA targeting *TP63* (2 sgRNA), *MED12* (3 sgRNA), non-targeting sgRNAs (2 sgRNA), or uninfected control T3M4 cells. -ΔΔCt values are plotted as the average of each sgRNA normalized to the average of housekeeping genes *ACTB* and *B2M*. For each gene perturbation, the average -ΔΔCt value is shown in a solid line, and the 95% confidence intervals are shown as translucid intervals. *MED12* knockdown inflection point (∼ day 5) is marked by a vertical black dashed line. **d,** RT-qPCR of *TP63* and its target genes *S100A2*, *IL1B* and *KRT6A* after *TP63* or empty vector cDNA overexpression and *MED12* (2 sgRNA) or non-targeting (2 sgRNA) CRISPRi sgRNAs. RNA was collected at day 5.5 post-infection with lentiviral sgRNAs, and three days after cDNA overexpression. mRNA fold change values are calculated as 2^-ΔΔCt^ normalized to the average of housekeeping genes *ACTB*, *B2M* and *PPIA*. *: double-sided t-test p-value<0.05. **e,** Metaplots and heatmaps of p63, MED12, and H3K27ac chromatin occupancy in KLM1 cells. Signal intensity values are centered around p63 peaks annotated by MACS2 and sorted by p63 intensity. **f,** Metaplots of p63 (left) and MED12 (right) genome occupancy signals centered around MED12 peaks at basal enhancers in KLM1 cells upon *ROSA26* or *TP63* knockout (2 sgRNA). **g,** ChIP-seq tracks of p63, MED12, and H3K27ac normalized occupancy at select basal-specific p63 direct target loci in KLM1 cells upon *ROSA26* or *TP63* knockout. Normalized enrichment values were generated with DeepTools bamCoverage -RPCG. **h,** Metaplots of p63 (left) and MED12 (right) genome occupancy signals centered around MED12-occupied peaks surrounding p63 direct target genes in SUIT2 CRISPRa-*TP63* lines. **e,f,h,** Metaplots and heatmaps were generated using DeepTools computeMatrix and plotHeatmap.

We next performed ChIP-seq analysis of p63 and MED12 localization across the genome of basal-like PDAC models. Chromatin occupancy profiling of p63 and MED12 revealed significant overlap in both KLM1 and T3M4 cells at active cis elements enriched for H3K27 acetylation (**Fig. 2e, Extended Fig. 2c**). This pattern of shared occupancy included a class of ‘basal lineage enhancers’ that we defined previously as selectively activated in basal-like PDAC models^18^. We hypothesized that p63, as a sequence-specific DNA-binding protein, was responsible for tethering MED12 to these basal enhancer elements. In support of this, genetic inactivation of p63 led to severe reductions of MED12 occupancy at basal lineage enhancers (**Fig. 2f,g, Supplementary Table 3**). In the converse experiment, we found that ectopic expression of p63 in a classical PDAC model was sufficient to acquire MED12 occupancy at core p63 targets in PDAC (**Fig. 2h, Extended Fig. 2d,e, Supplementary Table 3**). Taken together, these findings suggest that MED12 occupies basal lineage enhancer elements in a p63-dependent manner.

The findings above raised the possibility that p63 binds to the MED12-containing MKM to activate transcription of basal lineage genes. When transfected into human cells, we found that FLAG-tagged p63 immunoprecipitated endogenous MED12, as well as subunits of the core Mediator and the MKM (**Fig. 3a,b**). This association was not affected by the presence of ethidium bromide, suggesting it occurs in a DNA-independent manner (**Fig. 3b**). To evaluate this interaction in an alternative system, we expressed and purified recombinant MBP-tagged p63 from bacteria, which was competent for oligomerization and for sequence-specific DNA binding (**Extended Fig. 3a,b,c**). Pulldown experiments revealed that the MBP-p63 protein efficiently associated with endogenous MKM in nuclear lysates **(Fig. 3C**).

**Fig. 3.**
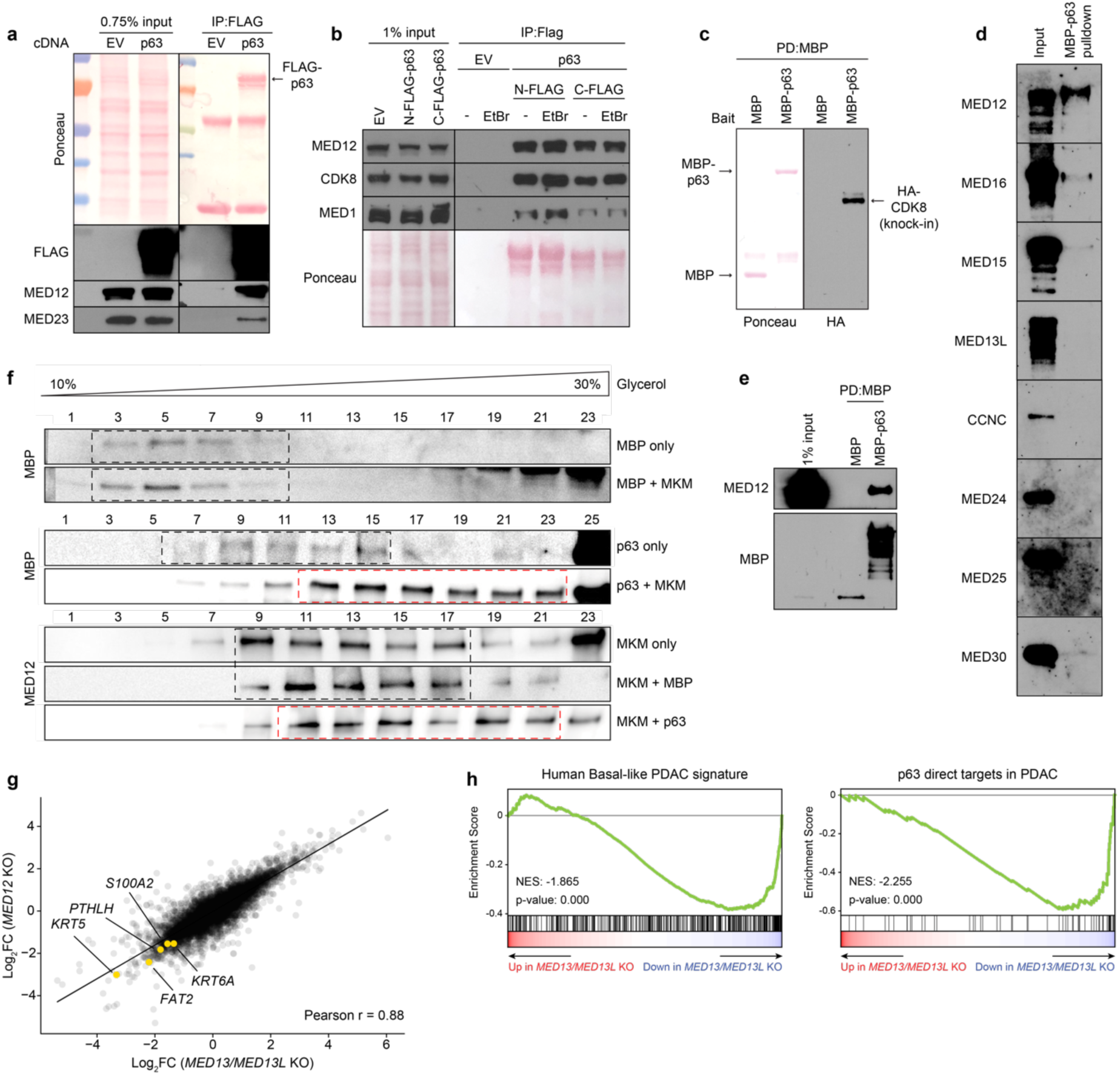
p63 directly binds to MED12 and the Mediator Kinase Module to activate the basal lineage program. **a,** FLAG immunoprecipitation of transiently transfected p63-C-3xFLAG in HEK293T nuclear lysates. Ponceau staining and western blot of FLAG and endogenous MKM (MED12) and core Mediator (MED23) subunits are shown for input and IP samples. The two bands in the Ponceau under p63 correspond to denatured IgG fragments. **b,** FLAG immunoprecipitation of transiently transfected N-3xFLAG-p63 or p63-C-3xFLAG in HEK293T nuclear lysates in the presence or absence of ethidium bromide (EtBr, final concentration of 50μg/mL). Ponceau and western blot of endogenous MKM (MED12, CDK8) and core Mediator (MED1) subunits are shown for input and IP samples. **c,** MBP-pulldown of purified MBP or MBP-p63 incubated with nuclear lysates of endogenously tagged 3NHA-CDK8 HeLa cells. Nuclear lysate was equally split between MBP and MBP-p63 for pulldown. Ponceau (left) shows the immobilized MBP fusion proteins, and HA stains the 3xHA-tagged CDK8. Only pulldown results are shown, as the signal was undetectable in the 1% input. **d,** Western blot of MBP-pulldown of purified MBP-p63 incubated with Sf9 cell lysates expressing different individual human Mediator subunits. 1% input is shown in the left lane. **e,** MBP pulldown of MBP or MBP-p63 incubated with human MED12-expressing Sf9 lysates. **f,** Glycerol gradient sedimentation of purified MBP or MBP-p63 incubated with human 4-subunit MKM. Glycerol percentages are indicated above the blots; larger complexes will migrate farther down the gradient. Black dashed boxes show glycerol gradient eluted fractions of individual p63, MBP, or MKM, as well as MBP-MKM incubation samples which did not display shifted migration pattern. Red dashed boxes highlight p63-MKM glycerol gradients that eluted in size-shifted fractions. **g,** Scatterplot depicting gene expression changes upon *MED12* or *MED13/MED13L* knockout. DESeq2-derived log2FC for all expressed genes are plotted. Representative plot of 2 different sgRNA per gene. Select basal genes are highlighted. **h,** GSEA plots of *MED13*/*MED13L* double knockout using gene signatures derived from human basal PDAC tumors (Chan-Seng-Yue et al. 2020, left) and direct p63 gene targets in PDAC (Somerville et al. 2018, right). Complete GSEA analysis for all the sgRNA tested can be found in Supplementary Table 2.

To evaluate potential direct interactions between p63 and Mediator proteins, we expressed nine different human MED subunits individually in Sf9 insect cells (which lacks endogenous human Mediator) and screened for an interaction with recombinant MBP-p63. Among the nine subunits tested, only MED12 bound efficiently to p63 (**Fig. 3d,e**). While this result suggested a direct p63:MED12 interaction, it is important to recognize that native MED12 exists as a stable subunit of the MKM. This prompted us to evaluate whether a p63:MKM complex can be reconstituted *in vitro*. We co-expressed human MED12, MED13, Cyclin C, and CDK8 in Sf9 cells and used size exclusion chromatography to isolate a stoichiometric MKM. This complex was incubated with MBP-p63 and evaluated using a glycerol gradient for a stable interaction. Notably, purified MBP-p63, but not MBP alone, was displaced towards heavier fractions of the glycerol gradient upon incubation with recombinant MKM (**Fig. 3f**). Together, these biochemical results suggested a direct interaction between p63 and the MED12-containing MKM.

Unexpectedly, we found that double *CDK8*/*CDK19* knockout or treatment with CDK8/CDK19 inhibitors failed to suppress p63 target gene expression in basal-like PDAC models despite leading to marked growth arrest (**Extended Fig. 3d, Supplementary Table 3**). This led us to hypothesize that the critical role of MED12 as an MKM subunit might be to bridge p63 with the core of Mediator, a function that is known to occur via its binding to MED13/MED13L^32^. While *MED13* and *MED13L* were each dispensable individually for basal lineage identity (**Fig. 1d**), the double knockout of these two paralogs led to highly similar transcriptional changes as the loss of *MED12*, including effects at basal lineage genes (**Fig. 3g,h, Extended Fig. 3e**). Of note, several core Mediator subunits also scored in our marker-based screen (**Fig. 1c**) and were found associated with p63 in nuclear lysates (**Fig. 3a,b**). Collectively, our genetic, epigenomic, and biochemical results support a mechanistic model in which MED12 and MED13/13L function as adaptors within the MKM that bridge p63 with the core of Mediator.

We next evaluated whether the growth of basal-like PDAC requires the interaction between MED12 and p63. To this end, we established a gene complementation assay evaluating whether different p63 cDNA constructs could rescue the growth-arrest caused by inactivation of endogenous *TP63* (**Fig. 4a**). Using competition-based cell fitness assays, we found that wild-type p63 and deletions of its sterile alpha motif (SAM) domain and transcription inhibition domain (TID) were still capable of supporting PDAC cell proliferation (**Fig. 4b**). In contrast, deletions of the DNA-binding domain (DBD) and the oligomerization domain (OD) of p63 behaved as null alleles, despite being expressed (**Fig. 4b, Extended Fig. 4a**). Guided by these results, we generated recombinant MBP-p63 ΔDBD and ΔOD proteins (**Extended Fig. 4b**). When compared to MBP-p63, both deletions abolished the interaction with MED12 in cell lysates (**Fig. 4c**). Taken together, these structure-function experiments suggested that the interaction between MED12 and p63 is required for the growth of basal-like PDAC cells.

**Fig. 4.**
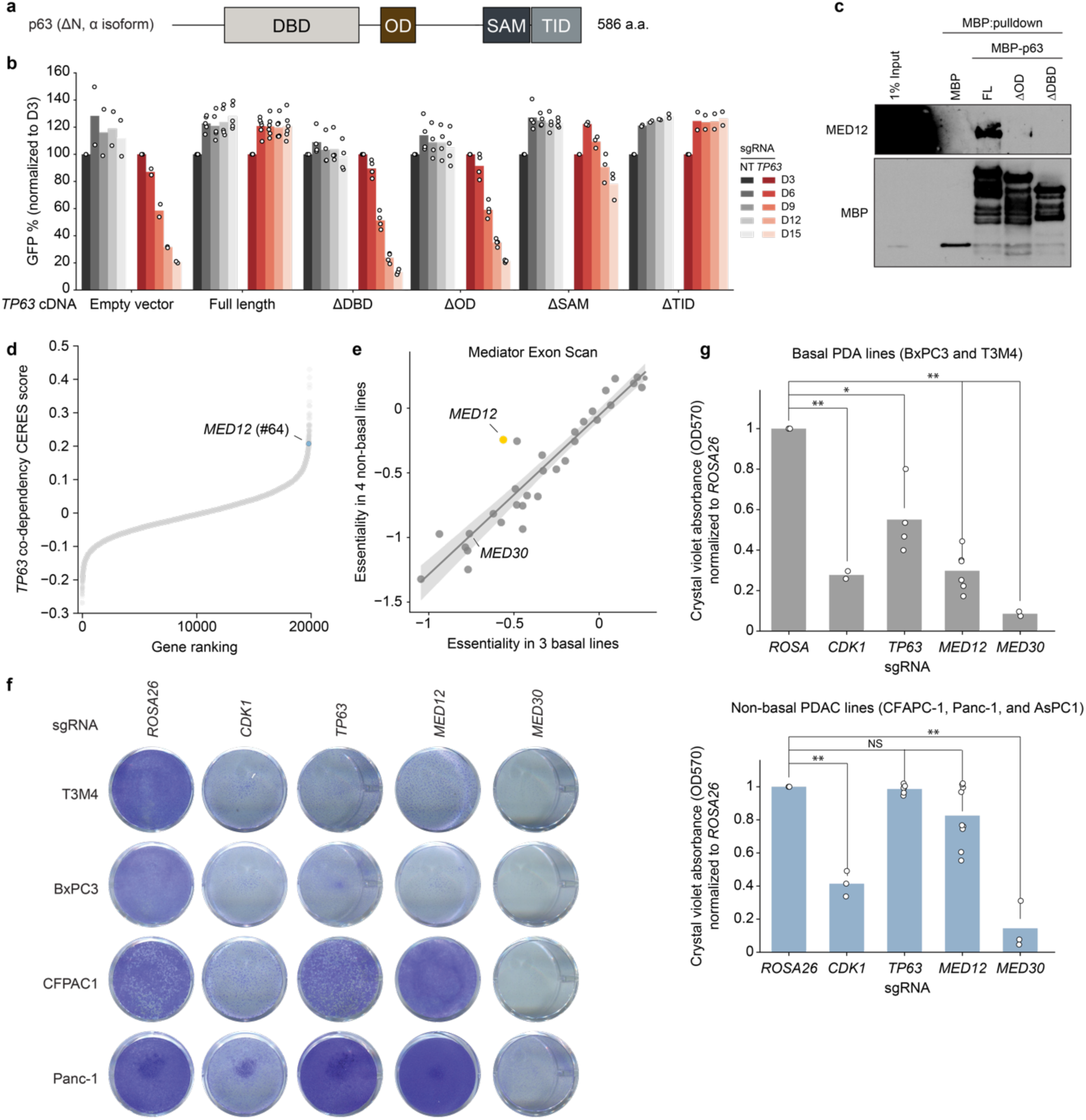
*MED12* is a lineage-biased genetic vulnerability of basal-like PDAC. **a,** Diagram of p63 (N-terminus ΔN, C-terminus α isoform) domain architecture. **b,** Gene complementation competition-based proliferation assay of different overexpressed p63 truncation mutants upon endogenous *TP63* CRISPRi-induced knockdown. **c,** MBP pulldown assay using purified MBP, full length MBP-ΔNp63 (FL), or MBP fusion p63 mutants lacking the oligomerization domain (ΔOD) or DNA-binding domain (ΔDBD). Purified protein was incubated with MED12-expressing Sf9 lysates overnight, followed by pulldown and western blot. **d,** Pearson r of >17.000 DepMap CERES gene codependencies with *TP63* across ∼1000 cell lines. **e,** Scatterplot of mean essentiality scores of all sgRNAs targeting each of 33 genes in the Mediator complex in basal (T3M4, BxPC3, and PK-1) and non-basal (SUIT2, AsPC-1, KLM1, and MiaPaCa-2) cell lines. Linear regression and 95% confidence interval are drawn. **f,** Crystal violet staining of basal (T3M4, BxPC3) or non-basal (CFPAC1, Panc-1) human PDA lines grown for 10 days after knockout of negative control (*ROSA26*), lethal control (*CDK1*), *TP63*, *MED12*, or panessential Mediator gene *MED30*. **b,** Quantification of re-solubilized crystal violet stain of basal (T3M4, BxPC3; top) or non-basal (CFPAC1, Panc-1, AsPC-1; bottom) grown for 10 days, as quantified by OD570. Two-sided t-test p-values: * < 0.05; ** <0.01; NS not significant.

To extend the findings above, we evaluated whether the growth of basal-like PDAC might be hypersensitive to genetic knockout of *MED12*. Remarkably, our analysis of the Cancer Dependency Map CRISPR screening data from >1,000 cancer cell lines^33^ revealed that *MED12* was in the top 1% of genetic co-dependencies of *TP63* among ∼18,600 genes that were evaluated (**Fig. 4d**), a correlation that did not exist for other core Mediator subunits (**Supplementary Table 1**). To evaluate this relationship more rigorously, we cloned a CRISPR tiling library that scanned 33 subunits of Mediator with all possible sgRNAs (∼10,000 sgRNAs in total), which we used to perform negative-selection CRISPR screens in three basal PDAC and four non-basal PDAC lines (**Supplementary Table 4**). While targeting of core Mediator subunits led to similar pattern of growth arrest in both groups of PDAC lines, our screens revealed that *MED12* was the Mediator subunit that had the strongest dependency bias towards basal-like PDAC (**Fig. 4e, Supplementary Table 4**). We validated this hypersensitivity using both Crystal Violet staining (**Fig. 4f,g**) and CellTiter-Glo proliferation assays (**Extended Fig. 4c**), demonstrating that basal-like PDAC lines were more dependent on *MED12* than classical PDAC despite sharing a similar dependency on core Mediator subunits.

Lineage plasticity is widely recognized as a phenotypic hallmark of human cancer that allows tumor cells to gain metastatic potential and evade therapy^1^. Recent advances in single cell transcriptomics and lineage tracing have provided unparalleled insights into this process^34,35^, however a major challenge exists in discovering perturbations that restrain cellular plasticity in human tumors. Building upon prior work evaluating smaller gene sets^18,20,23,36–38^, our study describes a genome-wide screening strategy that allows for the comprehensive mapping of genetic requirements to sustain an aberrant lineage state in PDAC. We anticipate that the methodology described here can be readily adapted to other clinically relevant tumor plasticity phenomena, such as adenocarcinomas that transition into neuroendocrine or mesenchymal cell states^2^. As many actionable targets for cancer therapeutics are ubiquitously expressed proteins across cell types, their involvement in lineage plasticity might only be revealed through high-throughput genetic perturbations. Our study also demonstrates how marker-based genetic screens can be leveraged to reveal a highly specific protein-protein interaction that functions as a lineage-biased cancer dependency. In this regard, the reconstituted interaction between p63 interaction MKM could be readily adapted into a biochemical assay for high-throughput chemical screening. Thus, our integrated experimental approach combining genetics and biochemistry may have biomedical utility to advance pharmacology that rewires cancer cell plasticity for therapeutic gain.

---

**Extended Fig. 1.**
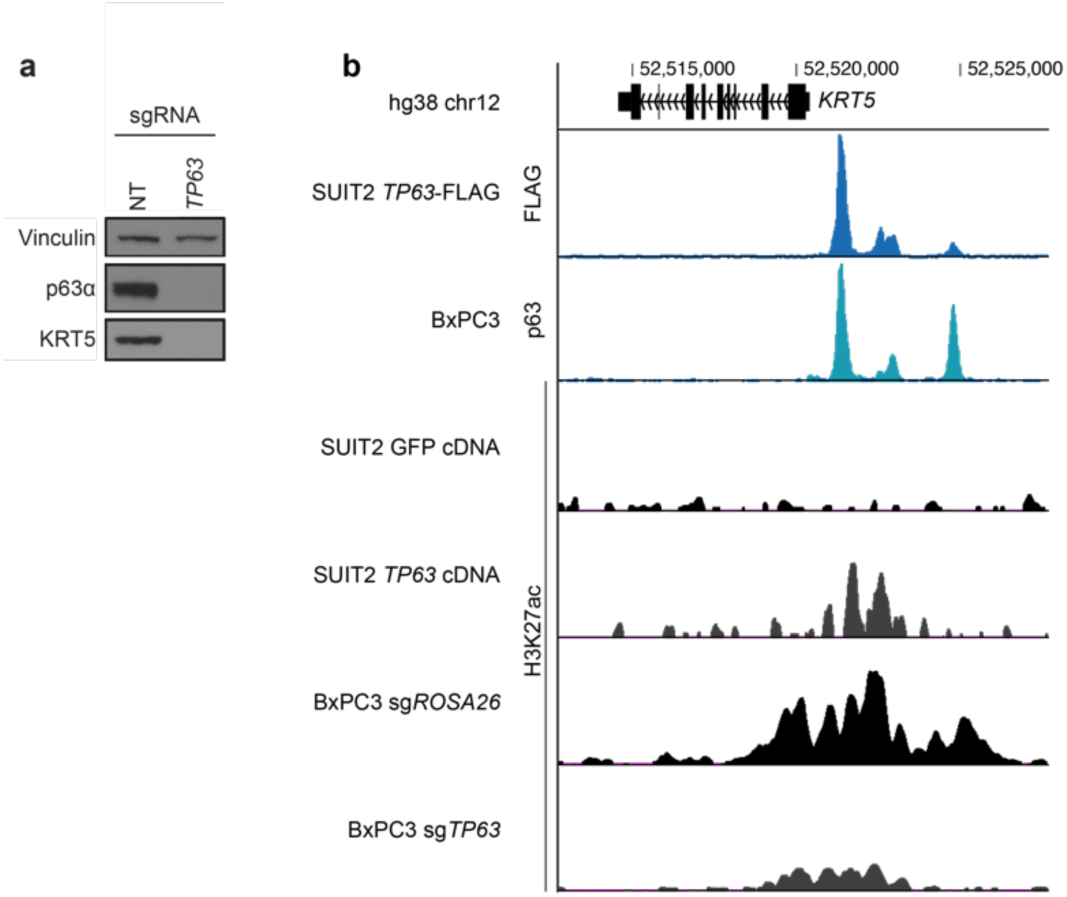
KRT5 expression is directly regulated by p63. **a,** Western blot of CRISPRi-mediated *TP63* or *KRT5* knockdown in KLM1 cells. **b,** ChIP-seq genomic occupancy tracks zoomed in the *KRT5* locus. The two upper tracks show normalized enrichment of endogenous (BxPC3) or overexpressed (p63-FLAG SUIT2) p63. The bottom four tracks show H3K27ac normalized enrichment after GFP or p63 overexpression in SUIT2 cells or *ROSA26* or *TP63* knockout in BxPC3 cells. ChIP signal was calculated using deepTools with the option BamCompare subtract to normalize each sample to its input. All tracks plotting ChIP data obtained with the same antibody are plotted in the same scale.

**Extended Fig. 2.**
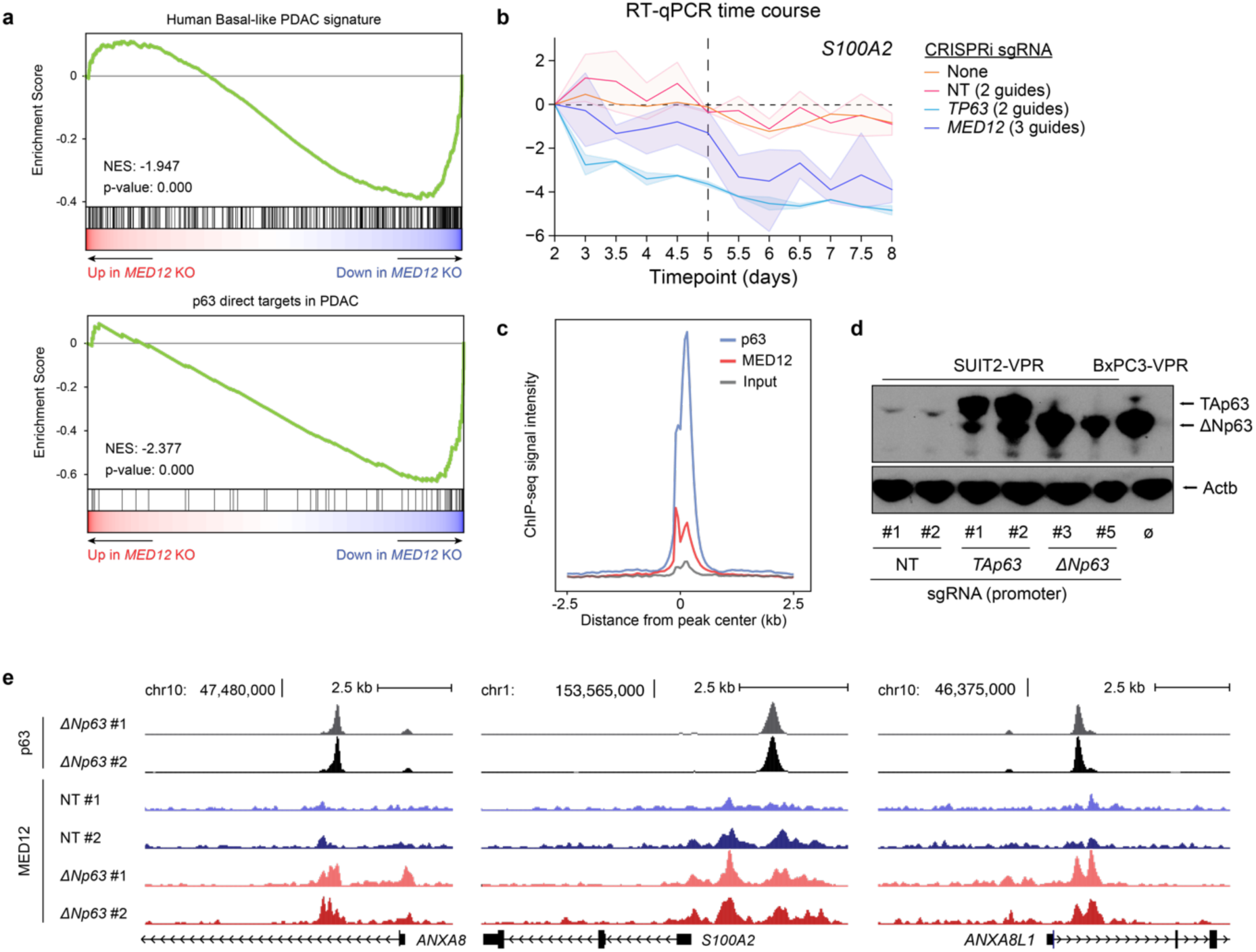
MED12 and p63 co-regulate the basal gene expression program. **a,** Representative GSEA plots of KLM1 *MED12* knockout using gene signatures derived from human basal PDAC tumors (Chan-Seng-Yue et al. 2020) and direct p63 gene targets in PDAC (Somerville et al. 2018). The plots were generated and the normalized enrichment scores (NES) and statistics were calculated using DESeq2-derived log2FC values of *MED12* vs *ROSA26* KO in GSEApy. Three biological replicates were used for each sample. Complete GSEA analysis for all the sgRNA tested can be found in Supplementary Table 2. **b,** Time-course RT-qPCR of *S100A2* after lentiviral infection with CRISPRi sgRNA targeting *TP63* (2 sgRNA), *MED12* (3 sgRNA), non-targeting sgRNAs (2 sgRNA), or uninfected control T3M4 cells. -ΔΔCt values are plotted as the average of each sgRNA normalized to the average of housekeeping genes *ACTB* and *B2M*. For each gene perturbation, the average -ΔΔCt value is shown in a solid line, and the 95% confidence intervals are shown as translucid intervals. *MED12* knockdown inflection point (∼ day 5) is marked by a vertical black dashed line. **c,** Metaplot of genomic occupancy of p63 and MED12 centered around p63 peaks in T3M4 cells. **d,** Western blot of SUIT2 CRISPR-activated *TP63* (isoform-specific) cells. BxPC3, which endogenously expresses the ΔN isoform of p63, is shown as a positive control in the rightmost lane. **e,** Genomic tracks of p63 and MED12 occupancy at direct p63 targets *ANXA8*, *S100A2*, and *ANXA8L1* in SUIT2-VPR lines infected with non-targeting (NT) or *TP63*-targeting sgRNAs.

**Extended Fig. 3.**
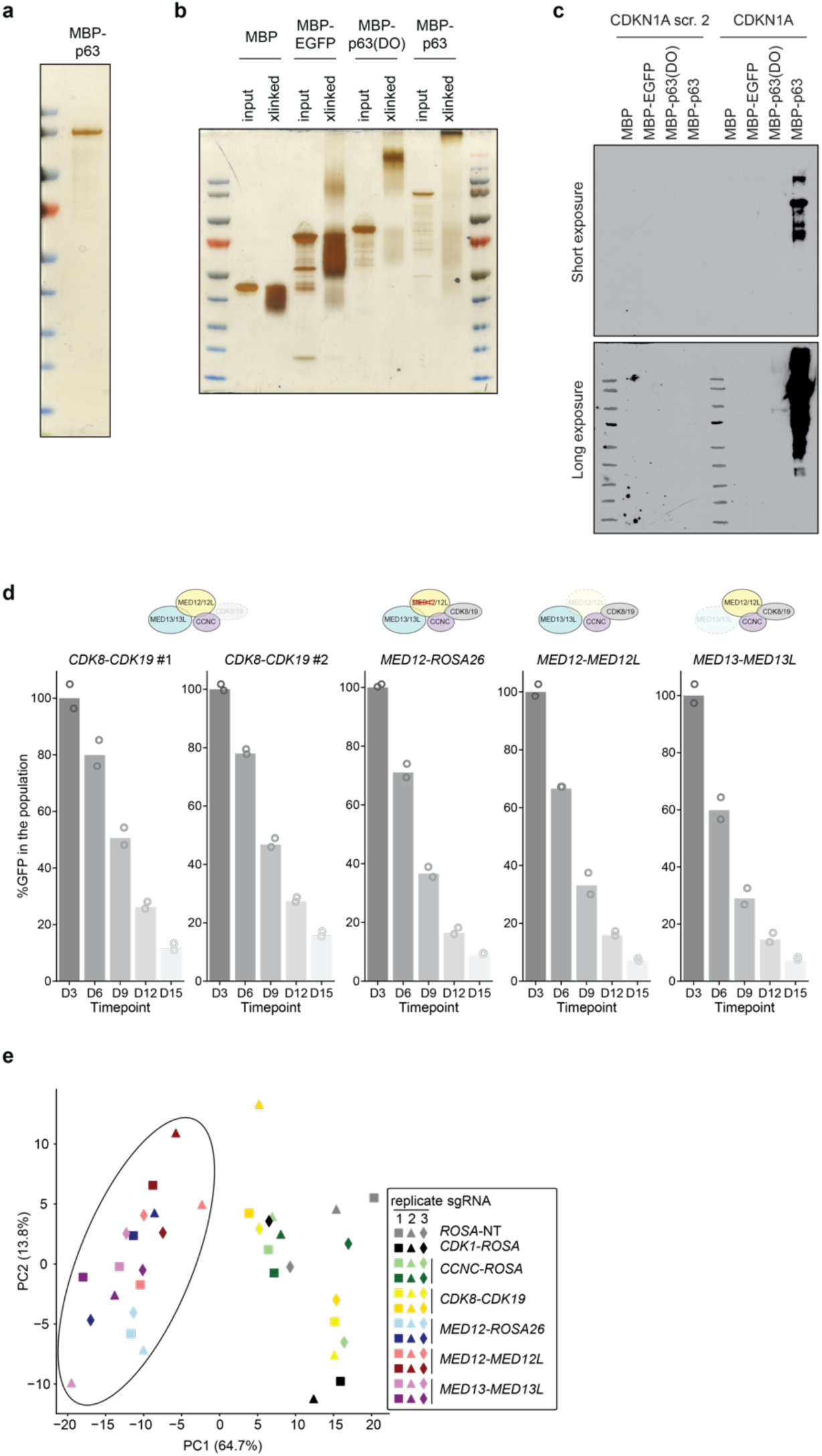
Characterization of recombinantly expressed and purified full length MBP-p63 protein and basal-related paralog redundancy within the MKM. **a,** Silver stain of full length purified MBP-p63. The single protein band was confirmed to be the expected MBP-p63 peptide by western blotting and mass spectrometry. **b,** Silver stain of 0.025% glutaraldehyde crosslinked (“xlinked”) and input purified full length MBP-p63, MBP-p63(DBD-OD), MBP-EGFP, or MBP alone. **c,** Western blot of DNA pulldown experiment using purified proteins and biotinylated DNA oligos containing the p63-binding sequence of the *CDKN1A* promoter or a scramble DNA control. **d,** Competition-based proliferation assays in Cas9-expressing T3M4 cells after lentiviral expression of the indicated sgRNA pairs (linked with GFP). Data are the mean normalized percentage of GFP (to day 3 after infection) of two to three sgRNAs. **e,** Principal component analysis of gene expression changes upon MKM paralog double knockout. *MED12*, *MED12*/*MED12L* and *MED13*/*MED13L* double knockouts are encircled together.

**Extended Fig. 4.**
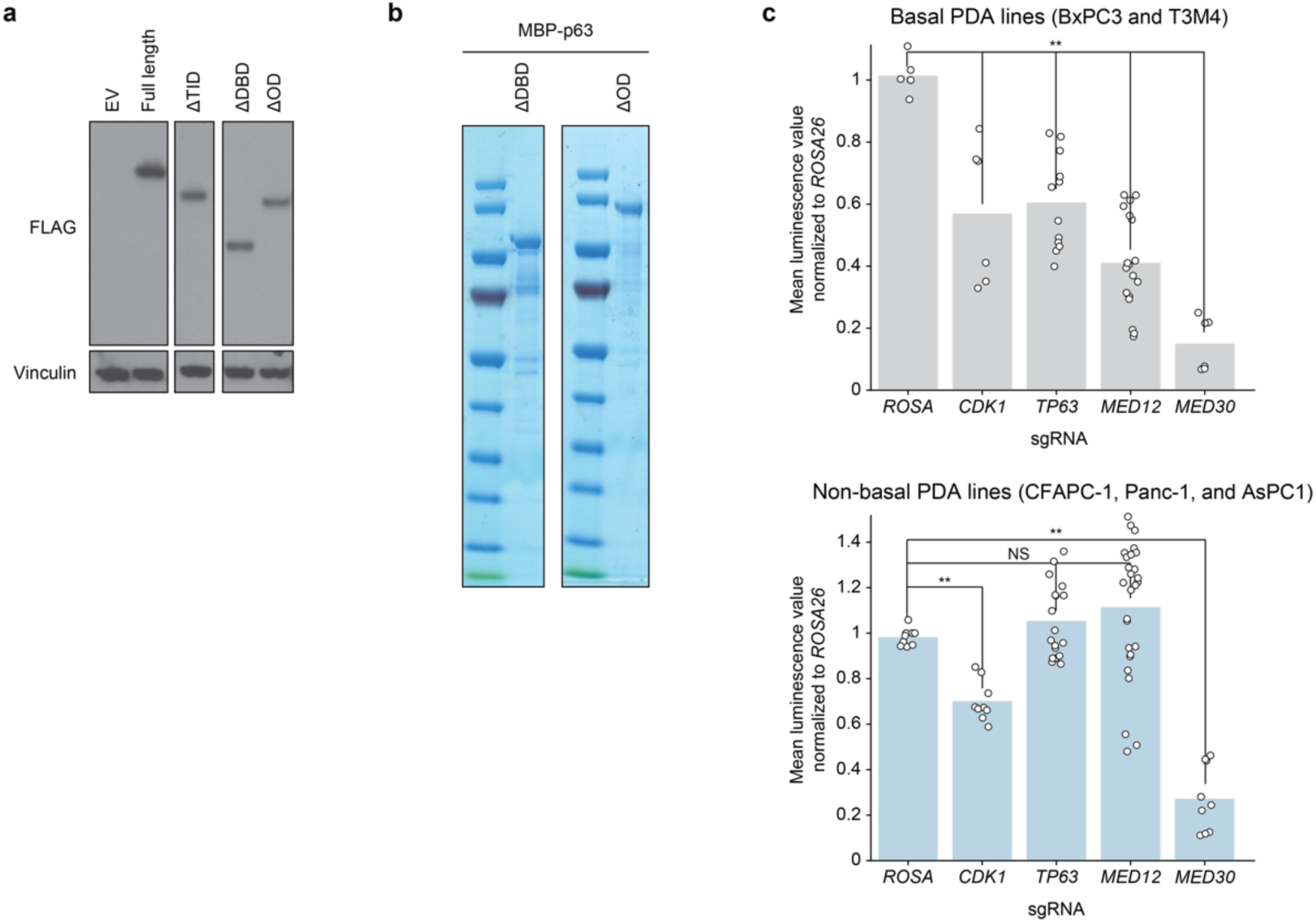
cDNA overexpression of p63 truncation constructs and CellTiter-Glo of *MED12* genetic targeting. **a,** Western blots of T3M4 cells stably expressing FLAG-tagged overexpressed truncated p63 cDNA used in gene complementation assays from Fig. 4b. **b,** Luminescence reading of CellTiter-Glo assay at day 8 post-infection with lentivirally-encoded sgRNA. All conditions were tested in triplicate. Two-sided t-test p-values: * < 0.05; ** <0.01; NS not significant.

## METHODS

### Human cancer cell lines and tissue culture

T3M4, KLM1, AsPC1, PK-1, and BxPC3 cells were cultured in RPMI supplemented with 10% FBS and penicillin/streptomycin (R10). SUIT2, CFPAC1, PANC1, MIAPaca2, and HeLa cells were cultured in DMEM supplemented with 10% FBS and penicillin/streptomycin (D10). Cell lines were purchased from commercial vendors and their identity validated by STR analysis. Cell lines were regularly tested for *Mycoplasma* contamination. All antibiotic concentrations used to select gene cassettes were empirically titrated in each cell line to achieve maximum selection with minimum toxicity.

### Plasmid construction and sgRNA cloning

The sgRNA lentiviral expression vector with optimized sgRNA scaffold backbone (LRG2.1T-puromycin, Addgene, #125594) was used for CRISPR knockout, interference and activation (a list of sgRNAs used in this study is provided in Supplementary Table 5). Stable cell lines were generated with the lentiviral Cas9-blasticidin vector (CRISPR knockout, Addgene, #125592), dCas9-KRAB-blasticidin (CRISPR interference, Addgene #89567), or dCas9-VPR-blasticidin (CRISPR activation, Addgene #63798 cloned into a lentiviral vector). *TP63* cDNA constructs were cloned into a lentiviral IRES-Neo vector using the In-Fusion cloning system (Takara). A 3×FLAG tag was added to *TP63* at the N- or C-terminus. MBP fusion *TP63* cassettes were cloned into a lac-inducible MBP-6xHis-TEV bacterial expression vector using In-Fusion cloning system (Takara).

### Lentiviral Production and Infection

Lentivirus was produced in HEK293T cells transfected with target plasmids and packaging plasmids (VSV-G and psPAX2) using polyethylenimine. Transfection media was replaced with fresh D10 8-12 hours following transfection, and lentivirus-containing supernatant was subsequently collected every 24 hours for 72 hours. All 2-3 collections were pooled and filtered through a 0.45μm filter. For lentiviral infections, cell suspensions or previously plated adherent cells were exposed to lentiviral-containing supernatant supplemented with polybrene to a final concentration of 4μg/ml. Lentiviral media was changed for fresh media after 12-48h.

### Intracellular FACS-based CRISPR Screens

CRISPR screens were conducted as previously described. Briefly, after determining the suitable lentiviral titer, ∼1-2×10^9^ Cas9-expressing cells were seed infected at D0 with genome-wide CRISPRi^28^ or knockout^31^ sgRNA library-encoding lentiviral suspension for a final 20%-30% GFP+ percentage of infected cells. Media was changed at 48h, and puromycin was added for 72h for sgRNA cassette integration selection. 6 days post infection, cells were resuspended in trypsin, counted, washed in cold PBS and fixed in -20C methanol at 10×10^6^ cells/mL under gentle vortexing. Cells were stored in methanol at -20C for a period up to 4 months. The day before sorting, cells were pelleted, washed 1x in FACS buffer (1% (w/v) ultrapure BSA, 0.5% (w/v) sodium azide, and 1mM EDTA in magnesium and calcium-free PBS), and incubated overnight in 1:400 primary antibody (KRT5 or p63) in FACS buffer at 10×10^6^ cells/mL rotating at 4C. The next day, cells were pelleted, washed 2x with FACS buffer, and incubated for 1h to 2h in 1:500 secondary antibody (AF647-conjugated anti-rabbit) in FACS buffer at 10×10^6^ cells/mL rotating at 4C protected from light. After washing 2x in FACS buffer, cells were resuspended in FACS buffer to ∼10×10^6^ cells/mL and sorted. Stained cells were sorted using a BD FACS Aria II cell sorter. The total number of cells sorted per screen was a minimum of 1500x the size of the sgRNA library. Cells were sorted into three different pools, with approximately 30% of cells sorted into the marker^low^ bin and 15% sorted into the marker^high^ bin. Cell pellets were then processed for DNA extraction and library preparation. A custom sequencing primer was added for next-generation sequencing of all CRISPRi screens as previously described^28^. All sequencing data from FACS-based genome-wide screens was analyzed with MAGeCK v0.5.9.3^39^ with the option MLE.

### Mediator exon scan library construction and cloning

sgRNA sequences targeting all 33 possible human Mediator subunits were retrieved from the CSHL in-house CRISPR sgRNA design tool, the VBC Score sgRNA database, and the Broad Institute CRISPick tool. Duplicate sgRNAs were removed, and all other remaining guides plus 391 non-targeting sgRNAs were incorporated into the final library pool (10.000 sgRNA).

Pooled amplicons with overhangs were ordered from Twist Bioscience, and cloned into the LRG-Puromycin (Addgene # 125594) backbone using NEBuilder HiFi DNA assembly master mix. The purified product was electroporated into MegaX DH10B (Invitrogen), and plated into LB-carbenicillin plates with a minimum representation of 100 colonies/sgRNA. The colonies were scraped from the plates and their DNA was extracted using Invitrogen MaxiPrep kit. Libraries were amplified for 10 cycles and sequenced to confirm successful cloning.

### Negative selection Mediator exon scanning CRISPR screens

∼1×10^8^ Cas9-expressing cells were seed infected with the Mediator exon scanning lentiviral sgRNA library to ∼20-30% by GFP+ population at day 3. At day 2 post-infection, ∼5×10^7^ cells were collected and flash frozen for early timepoint, and the remaining cells puromycin selected for 72h. Cells were subsequently passaged every three days for a period of 15 days, maintaining a minimum of ∼5×10^7^ cells in culture at any given time. After 15 days (late timepoint), ∼5×10^7^ cells were collected. Early and late timepoint cell pellets were treated with detergent and proteinase K, and its DNA extracted through phenol extraction. sgRNA depletion scores were calculated from raw sequencing files using MAGeCK v0.5.9.3^39^ with the option RRA. Downstream analysis was done in Python 3.6.

### DNA extraction and sgRNA sequencing for CRISPR screens

After sorting, cells from each sorting bin were pelleted and pooled, and their DNA extracted. Cells were resuspended in DNA extraction buffer (10mM Tris-HCl pH 8, 150mM NaCl, 10mM EDTA) at a density of 12.5M cells/mL. SDS and Proteinase K were added to final concentrations of 0.1% and 0.2mg/mL, respectively. The mixture was incubated for 48h at 56C, after which DNA was purified by phenol extraction. Equilibrated phenol was added 1:1 to the lysis mixture, mixed well, and centrifuged for 10min at 20,000 RCF. The supernatant was carefully removed, and another round of phenol purification was performed. DNA was then precipitated by adding 3 volumes of isopropanol and NaOAc pH 5.2 to a final concentration of 75mM and incubating overnight at - 20C. DNA was then pelleted at maximum speed for 1h, washed in 70% ethanol, and air-dried until translucent. After resuspension in water, DNA was assessed for quality by nanodrop before proceeding to library prep.

sgRNA were directly amplified from genomic DNA in one step PCR using NEBNext® Ultra™ II Q5® Master Mix (NEB). Each PCR reaction was done with 10µg of genomic DNA in 100µL final volume. Titrations of amplification cycles were performed for LRG (95C, 1min; n cycles [95C, 30s; 53C 30s; 72C 30s]; 72C 10min) and CRISPRi (98C, 30s; n cycles [98C, 10s; 65C 75s]; 65C 5min) sgRNA cassettes, which showed they could both be efficiently amplified using 22 cycles. All PCR reactions for each sample were pooled, and 400µL were taken for double-sided Ampure bead cleanup (0.65x + 1x bead volume) to preserve PCR amplicons (∼354bp and ∼274bp, respectively). Amplicons were sequenced using an Illumina NextSeq with 50% spike-in or pooled with high-diversity libraries (Cold Spring Harbor Genome Center, Woodbury).

### RNA extraction, RT-qPCR and RNA-sequencing

RNA Extraction and RT-PCR Total RNA was extracted using TRIzol reagent following the manufacturer’s instructions. 2 mg of total RNA was reverse transcribed to cDNA using qScript cDNA SuperMix, followed by RT-qPCR analysis with SYBR green PCR master mix on an ABI 7900HT fast real-time PCR system. Primers used for RT-qPCR are provided in Supplementary Table 5.

For RNA-seq experiments of CRISPR based targeting of *TP63*, *MED12*, and MKM paralogs, cells stably expressing Cas9 were infected with control or target sgRNAs in LRGP vector to >85% GFP positivity. RNA was collected at relevant timepoints as assessed by western blot kinetics of protein depletion and loss of cell viability. All gene knockout experiments were done with 3 biological replicates and 2-4 different sgRNA. Pharmacological inhibition experiments were done with 2 biological replicates.

RNA-seq libraries were constructed using the TruSeq sample Prep Kit V2 (Illumina) following the manufacturer’s instructions. Briefly, 2 mg of purified RNA was poly-A selected and fragmented with fragmentation enzyme mix. cDNA was synthesized with Super Script II master mix, followed by end repair, A-tailing, single-ended indexed adaptor ligation and PCR amplification. RNA-seq libraries were single-end sequenced for 76bp using an Illumina NextSeq platform (Cold Spring Harbor Genome Center, Woodbury).

### ChIP and ChIP-Seq Library Construction

For each ChIP reaction, cells were trypsinized, counted and resuspended to 5×10^6^ cells/mL. Cell suspensions were crosslinked in 1% formaldehyde at room temperature for 15 min, followed by addition of 0.125M glycine to quench the reaction. After two ice cold PBS washes, each 10 million cells were resuspended in 1mL of cell lysis buffer (10mM Tris pH8.0, 10mM NaCl, 0.2% NP-40) and incubated on ice for 7 min. After spinning down and decanting, nuclei were resuspended in 500μL of nuclei lysis buffer (50mM Tris pH8.0, 10mM EDTA, 1% SDS) and sonicated using a BioRuptor Pico water bath sonicator (30s on/off cycles, 1mL per tube). The number of cycles was adjusted to each cell line to obtain an average chromatin size distribution of 200-500 bp (T3M4: 16 cycles, KLM1: 17 cycles, SUIT2: 17 cycles). Each 500μL of sonicated chromatin from 10 million cells was diluted with 3.5ml of IP-Dilution buffer (20mM Tris pH 8.0, 2mM EDTA, 150mM NaCl, 1% Triton X-100, 0.01% SDS), and 1/20th of the sample was saved for input. Chromatin from 10 million (H3K27ac), 50 million (p63) or >=120 million (MED12) cells was incubated with 4μg (H3K27ac) or 10μg (p63, MED12) of the appropriate antibody and 50-200μL of magnetic protein A beads at 4°C overnight. Bead-bound chromatin was washed once with 1ml IP-wash 1 buffer (20mM Tris pH8.0, 2mM EDTA, 50mM NaCl, 1% Triton X-100, 0.1% SDS), twice with 1ml High-salt buffer (20mM Tris pH 8.0, 2mM EDTA, 500mM NaCl, 1% Triton X-100, 0.01% SDS), once with 1ml IP-wash 2 buffer (10mM Tris pH 8.0, 1mM EDTA 0.25M LiCl, 1% NP-40, 1% sodium deoxycholate), and once with 1ml TE buffer (10mM Tris-Cl, 1mM EDTA, pH 8.0). Chromatin was eluted from beads and reverse crosslinked in 400μL nuclei lysis buffer supplemented with 24μL of 5M NaCl and 1μg/mL RNase A by shaking (750rpm) overnight at 65°C. Protein digestion was done by adding 4μg/mL of proteinase K and incubating the mixture for 2 hours at 56°C. The DNA to be used for library prep was purified using QIAGEN PCR purification kit and following manufacturer’s instructions.

ChIP-seq library was constructed using Illumina TruSeq ChIP Sample Prep kit following the manufacturer’s protocol. Briefly, ChIP DNA was end repaired, A-tailed and ligated to Illumina compatible single-indexed adaptors. 15-18 PCR cycles were used for final library amplification, which was purified 2x with 1.1x Ampure XP beads and subsequently analyzed on a Bioanalyzer using a high sensitivity DNA chip (Agilent). ChIP-seq libraries were single-end sequenced for 76bp using an Illumina NextSeq platform (Cold Spring Harbor Genome Center, Woodbury).

### General computational and statistical analysis

All sequencing data was analyzed using CSHL high performance computing system. Packages used to analyze next generation sequencing data were installed in independent Anaconda environments to minimize conflicts of dependencies. Downstream analysis was performed using Python 3 in JupyterLab notebooks. Statistical tests and regressions were done using Scipy and Seaborn.

### RNA-Seq Data Analysis

Single end 76bp raw sequencing reads were mapped to the hg38 genome using STAR v2.7.9a^40^, followed by HTSeq v2.21^41^ to generate raw gene counts. Low abundance transcripts were filtered out (<2 average raw counts), and normalized transcripts (variant stabilized transcripts) and differential expression values were calculated using DESeq2^42^. Three biological replicates were used per sgRNA per cell line tested.

Gene set enrichment analysis was conducted using GSEApy^43^. GSEA was performed using variant stabilized transformed counts of each tested sample in duplicates or triplicates and the permutation ‘gene set’ option. Gene signatures for human basal PDAC was retrieved from signature 10 of Chan-Seng-Yue et al. 2020, and for direct p63 target genes in PDAC from Somerville et al. 2020. GSEA normalized enrichment scores and p-values calculated for all RNA-seq samples of this study, as well as the list of genes in each signature used, are displayed in Supplementary Table 2.

### ChIP-Seq Analysis

Single end 76bp sequencing reads were mapped to the hg38 genome using Bowtie2 with default settings^44^. MACS v2.2.6 ^45^ was used to call peaks using input genomic DNA as control. Annotation of ChIP-seq peaks was performed using HOMER v4.11 with default settings^46^. To visualize genomic tracks, bigWig files were generated from BAM files using deepTools v3.5.0 bamCoverage function normalizing with reads per genome coverage^47^. To define BED files of peaks and peak overlaps, MACS2 output narrowPeak or broadPeak files were merged using bedtools v2.30.0 merge and intersect tools^48^. Heatmaps and average chromatin occupancy metaplots were generated using computeMatrix and plotHeatmap functions of deepTools, taking bigWig files and BED files as input.

For ChIP-seq analysis of KLM1 *TP63* KO, MED12 peaks were merged between all KLM1 samples and intersected with a list of basal lineage enhancers retrieved from Somerville et al. 2018 using bedtools. The resulting 453 loci were evaluated for MED12 and p63 abundance upon *ROSA26* or *TP63* KO using Deeptools.

For SUIT2 CRISPRa *TP63*, MED12 peaks were merged between all SUIT2 samples and annotated using HOMER. 80 MED12 peaks whose nearest genes intersect with p63 direct target genes from Somerville et al. 2018 were used to generate metaplots of all SUIT2 CRISPRa conditions centered around these loci. The custom cis-regulatory elements defined in this study are shown in Supplementary Table 3.

### CRISPR-Based Targeting for competition assays

For GFP-depletion assays, GFP% was measured using a MACSQuant® Analyzer Flow Cytometer on day three post viral transduction (P0) with lentivirally-encoded sgRNA, and then every three days until the end of the experiment. All competition data was collected in duplicates or triplicates.

### Gene complementation assays

*TP63* wildtype and mutant cDNA were stably expressed in T3M4 cells, which were selected with G418 for 7 days to generate stable populations (confirmed by FLAG western blot). At day 0 of the assay, cells were infected with lentiviral CRISPRi sgRNA targeting *TP63* (2 sgRNA) or non-targeting (2 sgRNA) to ∼50% BFP^+^ rates. The BFP^+^ proportion of the population was followed every three days until day 15 post-infection using a MACSQuant® Analyzer Flow Cytometer.

### Crystal violet assays

For crystal violet assays, 40.000 Cas9-expressing cells were plated in 12 well plates and spinfected 24h later with lentiviral sgRNAs. After 48h, cells were trypsinized and re-seeded. At day 10 post-infection cells were washed 1x with 1mL of ice-cold PBS and then fixed in 1mL of methanol for 10 minutes at -20C. After removing the methanol, 1mL of 0.5% crystal violet solution dissolved in 20% methanol was added, and cells were stained by gently rocking the plates for 15 minutes at RT. The stain was decanted, and plates were washed 3x with streaming water to remove background staining. Plates were dried overnight and scanned. To quantify the signal, the crystal violet staining was solubilized with 300μL of 30% acetic acid by gently rocking for 15 minutes at RT. 200μL were transferred to a 96 well plate and its OD (570 nm) was acquired using a Molecular Devices SpectraMax i3.

### CellTiter-Glo

Cells were grown in opaque white 96 well plates for the duration of the assay. At endpoint, CellTiter-Glo reagent was added at a buffer:media ratio of 1:2, and incubated at RT for 10 min on a rocker. Luminescence was subsequently measured using a Molecular Devices SpectraMax i3.

### Bacterial expression and purification of recombinant MBP-p63

MBP-6xHis-TEV-p63 expression was induced in BL21 cells grown in Luria broth supplemented with antibiotics and 0.5mM IPTG for 18h at 16C. Cells were collected by centrifugation at 4000 RCF for 10min at 4C, and resuspended in 30 mL lysis buffer (50mM Tris pH 7.5, 150mM KCl, 5mM MgCl_2_, 0.05% NP-40, 10% glycerol, 0.2mM EDTA, 1mM DTT (fresh)) supplemented with protease inhibitors and 100μg/mL lysozyme. After 45min incubating on ice, cells were sonicated for 2 minutes and 30 seconds (3s on, 5s off) with a probe sonicator. Lysates were clarified by ultracentrifugation at 100,000 RCF for 1 hour at 4C. Cleared bacterial lysate from 1L of culture was incubated with 2mL of equilibrated amylose resin (NEB #E8021L) in a rotator overnight at 4C. The next day, the resin was pelleted, washed 1x in 50mL wash buffer (50mM Tris pH 7.5, 500mM KCl, 5mM MgCl_2_, 0.05% NP-40, 10% glycerol, 1mM DTT (fresh)) and resuspended in 15mL lysis buffer. After passing the lysate and resin mix through a chromatography column, the protein was eluted in 5 fractions of 1mL of elution buffer (20mM maltose in lysis buffer). A size exclusion purification step followed using an AKTA Pure 25M (Cytiva 29018226) using a Superdex 200 Increase 10/300 GL. Purity was evaluated by SDS-PAGE and Coomassie staining, and protein was flash frozen in liquid nitrogen and kept at -80C. All purified proteins were validated by mass spectrometry peptide identification and western blot.

### Sf9 expression and purification of recombinant human MKM and other Mediator genes

Each individual Mediator subunit was cloned and expressed in a pLIB plasmid (Addgene #80610). *MED12*, *MED13L*, Twin-Strep-*CDK8*, and *CCNC* were cloned into a multiBac vector subsequently used to generate bacmid with the EmBacY vector (Geneva Biotech). To generate baculovirus, Sf9 cells were transfected with TransIT-insect transfection reagent (Mirus) and 5-10μg of plasmid following manufacturer’s instructions. A baculovirus amplification round was then performed with this supernatant using 200mL of Sf9 in suspension culture seeded at 10^6^/mL and grown for 48-72h. 20-50mL of baculovirus was used to infect 1L of Sf9 for protein expression. The cultures were grown for 48-72h, at which point cells were collected by centrifugation. Pellets were kept frozen at -80C or proceeded immediately for protein extraction. For protein isolation, pellets were resuspended in lysis buffer (25mM HEPES pH 7.5, 100mM KCl, 0.05% NP-40, 10% glycerol) and Dounce homogeneized 40 times on ice using pestle A. Debris was removed by ultracentrifugation at 100.000g for 1h at 4C, and the supernatant was either flash frozen or subjected to further purification. For MKM purification, twin-strep tagged MKM containing lysates were incubated with Streptactin XT magnetic bead slurry (IBA Lifesciences) and purified according to manufacturer’s recommendation. Further purification using a glycerol gradient was performed before experiments where high purity was required. Protein preparations were evaluated by SDS-PAGE and Coomassie or silver staining, and protein was flash frozen in liquid nitrogen and kept at -80C.

### Glycerol gradients

Glycerol gradients were prepared using a peristaltic pump and pre-cooled base buffers (10-30% glycerol, 20mM 7.6 HEPES-HCl, 150mM KCl, 0.02% NP-40, 0.1mM EDTA). Lysates and/or purified proteins in a maximum volume of 500µL were incubated for 3h before immunoprecipitating and carefully layering the elutions on top of a freshly poured 2.4mL glycerol gradient. Samples were ultracentrifuged in a TLS 55 rotor at 35,000 RPM for 16h at 4C, with the standard acceleration and deceleration speed settings. 100 µL fractions were successively taken from the top layer without disrupting the column, evaluated by Coomassie, silver stain and/or western blot, and taken to further processing and experimenting.

### Immunoprecipitation

HEK293T cells were collected by trypsinization or scraping, and washed 1x with ice-cold PBS. Cells were then incubated with 2mL of cell lysis buffer (25mM HEPES pH 7.5, 10mM KCl, 1.5mM MgCl_2_, 0.05% NP-40, 15% glycerol, 1mM DTT (fresh), protease inhibitors (fresh)) per 10cm dish of cells for 10min. Nuclei were then pelleted for 5min at 3000RCF, and incubated in nuclear extraction buffer (25mM HEPES pH 7.5, 150mM KCl, 1.5mM MgCl_2_, 0.05% NP-40, 2 mM HEPES pH 7.5) for 30min. Nuclear lysates were cleared by centrifugation at 20.000RCF for 1h, and subsequently incubated with magnetic M2 FLAG beads overnight on a rotator. After washing 4x with 1mL of nuclear extraction buffer, beads were resuspended in 50µL of nuclear extraction buffer. After adding 50µL of 2x Laemmli buffer, samples were boiled for 10min at 98C, and ran on an SDS-PAGE gel for western blotting. Except for the sample boiling, all steps were performed at 4C.

### MBP pulldown

50-100 µL of magnetic amylose bead slurry was equilibrated in 20-40 volumes of lysis buffer (20mM HEPES pH 7.5, 150mM KCl, 0.05% NP-40%), and incubated with MBP or MBP-tagged protein for 2-3h. After washing 1x with 10-20 volumes of lysis buffer, the protein-resin mix was incubated with pre-cleared nuclear lysates or purified protein overnight. After washing 4x with 10-20 volumes of lysis buffer, the resin was resuspended in 100 µL of lysis buffer supplemented with 20 mM of maltose. The mixture was incubated for 30 min on a nutator, after which the eluate was separated from the resin and boiled at 98C for 5 min in 1x Laemmli buffer. Except for the sample boiling, all steps in the protocol were performed at 4C.

### Protein crosslinking

Purified protein solutions were incubated with 0.025% of glutaraldehyde on a nutator for 30min at 4C. After crosslinking, samples were directly boiled in 1x Laemmli buffer at 98C for 10min and loaded on an SDS-PAGE gel.

### DNA pulldown

5′ biotinylated single-stranded DNA oligos (IDT) were generated hybridized with complementary oligonucleotides in a 1:1 ratio in duplex buffer (IDT) by heating at 95 °C for 5 min followed by slow cooling to 25C at 5 °C min^−1^ in a thermocycler. For each DNA pulldown, 25 µl of Dynabeads MyOne Streptavidin T1 beads (Thermo Fisher Scientific, 65602) were washed with binding buffer (20mM Tris (pH 7.5), 1M NaCl, 1mM EDTA) followed by two washes with PBS. DNA probe (5pmol) was incubated with 25µl washed MyOne T1 beads (Thermo Fisher Scientific, 65001) in 500μl of binding buffer at 4C with rotation for 1 h, followed by two washes with binding buffer, one wash with PBS and two washes with NTN150 buffer (20mM Tris (pH 7.5), 150mM NaCl, 0.5% NP-40). DNA on beads was then incubated with purified protein 2h incubation. Protein-bound T1 beads were washed twice with NTN150 buffer and once with PBS. Proteins were then eluted from beads by heating at 98C for 10 min in 200µl Laemmli sample buffer with 5% β-mercaptoethanol. After 5min of centrifugation at maximum speed in a tabletop centrifuge, samples were analyzed by western blotting.

### Western blots and protein analysis

Cells were counted in each sample and normalized for cell number. Equal number of cells was then washed 1x in PBS, and resuspended in 1x Laemmli buffer diluted in PBS with 5% 5% β-mercaptoethanol. Samples were boiled at 98C for 5min, and stored at room temperature for a maximum of one month. Just before loading on an SDS-PAGE gel, samples were boiled for 3min and centrifuged at 20.000RCF for 5min.

### Mediator knock-in cell line generation and validation

HeLa S3 were grown in adherent tissue culture conditions in D10. To knock-in a triple HA cassette in the N-terminus of the CDK8 locus, cells were trypsinized, counted and washed 1x in PBS. 1.2×10^6^ cells were pelleted and resuspended in RT Lonza SE buffer. 180 pmol of Cas9 protein (IDT) were complexed with 360 pmol of sgRNA (IDT) previously for 10 minutes at RT. The Cas9-sgRNA complex was added to the cells, along with ∼2-4μg of double stranded HDR template cassette at ∼2μg/μL containing the sequences to knock-in (hygromycin-resistance gene - IRES - 3xHA) flanked by 50bp sequences. The mixture was then electroporated according to manufacturing instructions, and rapidly seeded in pre-warmed media. Cells were left to recover for 3 days, hygromycin selected for 5 days, and single cell cloned by flow cytometry-assisted sorting. Clones were expanded and screened for expression of tagged target protein and the ability of the tagged protein to co-immunoprecipitate the Mediator complex by western blot and mass spectrometry.

### Antibodies and protein reagents

All antibodies and protein-binding matrices, as well as their concentrations for each application, can be found in Supplementary Table 6.

### Data availability

All genomic datasets are available at the GEO database under accession code GSE229062 (reviewer token: mlgfqgqgpvstzah). KLM1 H3K27ac ChIP-seq data was obtained from Somerville et al. 2020. The cancer dependency dataset as well as expression was obtained online (https://depmap.org/portal/download/, DepMap Public 21Q4).

## ACKNOWLEDGEMENTS

We acknowledge B.W. Stillman, M.G. Hammell and J.M. Sheltzer for discussions and suggestions throughout the course of this study. This work was supported by Cold Spring Harbor Laboratory NCI Cancer Center Support grant CA045508. Additional funding was provided to C.R.V. by the Pershing Square Sohn Cancer Research Alliance, National Institutes of Health grants CA013106 and CA229699, and the Cold Spring Harbor Laboratory and Northwell Health Affiliation. D.M.-S. was supported by a Boehringer Ingelheim Fonds Ph.D. fellowship. D.J.T. was supported by R35 GM139550 and A.C.S. was supported in part by T32 GM008759.

## AUTHOR CONTRIBUTIONS

D.M.-S. and C.R.V. conceived this project, designed the experiments and wrote the manuscript with input from all of the authors. D.M.-S. performed the experiments and analyzed the data with the following help: A.C.S. isolated MKM and performed glycerol gradients in figure 3f; D.S. conducted cell culture and cell sorting for basal lineage CRISPR screens; V.K. performed knockdown RNA-seq experiments; A.A. and J.L. helped establish systems for Mediator complex expression in Sf9; C.R.V. and D.J.T. supervised the studies. C.R.V. acquired the funding.

## SUPPLEMENTARY INFORMATION

Supplementary Table 1. Marker-based CRISPR screening results. Supplementary Table 2. Gene signatures and GSEA results.

Supplementary Table 3. Basal lineage loci from ChIP-seq.

Supplementary Table 4. Mediator Exon Scan library and CRISPR screen results. Supplementary Table 5. sgRNA and RT-qPCR primer sequences.

Supplementary Table 6. Antibodies and reagents.

